# CellSwarm: LLM-Driven Cell Agents Recapitulate Tumor Microenvironment Dynamics and Sense Indirect Genetic Perturbations

**DOI:** 10.64898/2026.02.25.707926

**Authors:** Xuanlin Meng, Tengjiao Wang, Zhongyuan Dong, Xinhao Li, Xiuliang Cui, Lianghua Wang

## Abstract

Agent-based models of the tumor microenvironment (TME) traditionally rely on hand-coded rules that cannot generalize beyond their programmed logic. Here we present CELLSWARM, a framework that replaces rule-based cell decision-making with large language model (LLM)-driven autonomous agents. Each simulated cell maintains persistent state, 14 signal pathways, and a memory stream, with an LLM serving as its cognitive core. Using structured knowledge bases for cancer-specific context, CELLSWARM recapitulates TNBC microenvironment composition with fidelity comparable to hand-coded rules (Jensen–Shannon divergence 0.144 vs. 0.146; P=0.012 vs. random, Mann–Whitney U test). Beyond matching rule-based performance, LLM-driven agents demonstrate three capabilities absent from rule-based models: cross-cancer generalization by swapping knowledge base entries, treatment response prediction concordant with clinical data (anti-PD-1: 17.6% simulated vs. 21% clinical), and sensing of indirect genetic perturbations that propagate through intermediate signaling cascades (IFN-γ KO: Agent +15.7% vs. Rules +0.3%; P=0.005). CELLSWARM demonstrates that LLM-driven cell agents can recapitulate and extend TME simulation beyond the reach of hand-coded rules.

## Introduction

Tumors are not isolated masses of malignant cells but dynamic ecosystems composed of tumor cells, immune cells, stromal cells, and vascular and lymphatic networks. The classical hallmarks framework emphasized the systemic, multi-stage nature of acquired tumor capabilities, and subsequent revisions incorporated immune evasion, inflammation, metabolic reprogramming, and microenvironment remodeling as additional dimensions, reinforcing the view that tumor progression and treatment response depend critically on microenvironmental coupling (Hanahan and Weinberg 2000, 2011). Research centered on the tumor microenvironment (TME) has further established that cellular composition, spatial architecture, diffusion gradients, and signaling interaction networks collectively determine tumor evolutionary trajectories and therapeutic accessibility (Quail and Joyce 2017). The “immune contexture”—encompassing immune cell types, functional states, and spatial organization—correlates tightly with clinical outcome and varies substantially across tumor types and individuals (Fridman et al. 2012).

Immune checkpoint blockade (ICB), which unleashes anti-tumor immunity by relieving inhibitory pathway engagement, has produced breakthrough responses across multiple cancer types (Pardoll 2012). Yet both clinical trials and real-world evidence show that the fraction of patients who benefit remains limited: the immune-inflamed, immune-excluded, and immune-desert subtypes of the tumor immune microenvironment dictate sensitivity and resistance to immunotherapy (Binnewies et al. 2018). Key translational challenges include the scarcity of transferable preclinical models, insufficient mechanistic interpretability, a combinatorial treatment space too vast to search systematically, and reproducibility problems driven by inter-patient heterogeneity (Hegde and Chen 2020).

Triple-negative breast cancer (TNBC) illustrates these difficulties. Although PD-1/PD-L1 inhibitors show clinical activity in selected subgroups, overall response rates remain modest and are strongly dependent on baseline immune status and microenvironment structure (Nanda et al. 2016; Adams et al. 2019). Suppressive mechanisms in the TME are typically multifactorial; metabolic competition and nutrient/oxygen limitation are important drivers of effector T cell exhaustion and decline of key effector molecules such as IFN-γ (Chang et al. 2015). Meanwhile, large-scale TME subtyping studies suggest that relatively conserved microenvironment subtypes exist across cancer types and can predict immunotherapy response, but how these statistical associations map onto executable cell-level behavioral rules—and how to perform mechanistic reasoning and counterfactual testing at the individual level—remains without a unifying framework (Bagaev et al. 2021).

Single-cell transcriptomics and spatial omics have sharply increased the resolution at which we can dissect the TME, enabling characterization of tumor ecosystems at the level of cell types, states, and interactions, and uncovering complex crosstalk networks between tumor cells and immune/stromal compartments (Tirosh et al. 2016). In solid tumors such as breast cancer, single-cell and spatially resolved data resources now systematically capture immune infiltration patterns, tumor cell state lineages, and their spatial organization, providing a data foundation for individualized mechanistic modeling (Wu et al. 2021). Multi-modal microenvironment studies have further shown immune cell remodeling and spatial heterogeneity at the tissue scale, emphasizing that spatial–functional coupling is necessary for reliable inference of treatment response (Klemm et al. 2020).

On the modeling side, multiscale and agent-based tumor simulations can explicitly represent individual cells, local interactions, and diffusion/mechanical constraints, supporting interpretable “rule-to-phenotype” reasoning and enabling exploration of treatment strategies, dosing regimens, and combination therapy spaces (Wang et al. 2015). Open-source platforms such as PhysiCell provide efficient simulation kernels for 3D multicellular systems with extensible phenotype modules, lowering the engineering barrier to building reusable models (Ghaffarizadeh et al. 2018). However, current agent-based models remain heavily dependent on researcher-specified rules, parameters, and interaction structures. When transferring across datasets or patients, rule extraction, parameter calibration, and uncertainty characterization become bottlenecks that limit application to large-scale individualized inference and automated model discovery.

In parallel, the rapid advances in large language models (LLMs) have opened new approaches to knowledge organization, rule generation, and tool invocation through natural language interfaces. Models exemplified by GPT-4 show strong general capabilities in multi-task reasoning, code generation, and complex instruction following (OpenAI et al. 2023). Retrieval-augmented generation (RAG) couples external knowledge bases with generative models, providing a path toward traceable factual grounding and dynamic knowledge updates (Lewis et al. 2020). The ReAct paradigm further interleaves reasoning traces with executable actions, supporting iterative planning and error correction through environment interaction (Yao et al. 2022). Related work on generative agents has shown that LLMs combined with memory, reflection, and planning modules can produce credible multi-agent behaviors and emergent phenomena in complex settings (Park et al. 2023).

Building on these advances, we propose CELLSWARM, an LLM-driven multi-agent modeling framework for the tumor immune microenvironment. CELLSWARM organizes single-cell and spatial data together with biological priors—signaling pathways, cell–cell interactions, and metabolic constraints—into retrievable knowledge bases, and during simulation the LLM performs rule selection, parameter suggestion, and experimental design within an interpretable mechanistic world. This enables inference of immunotherapy response, counterfactual testing of key mechanisms, and systematic search over candidate intervention strategies. We validate CELLSWARM across six cancer types spanning the immunological spectrum (Fig. 2–Fig. 3), show that it senses indirect genetic perturbations that rule-based models cannot detect (Fig. 4), establish robustness across multiple LLM backends (Fig. 5), and dissect the mechanistic basis of emergent simulation dynamics (Fig. 6).

**Figure. 1.**
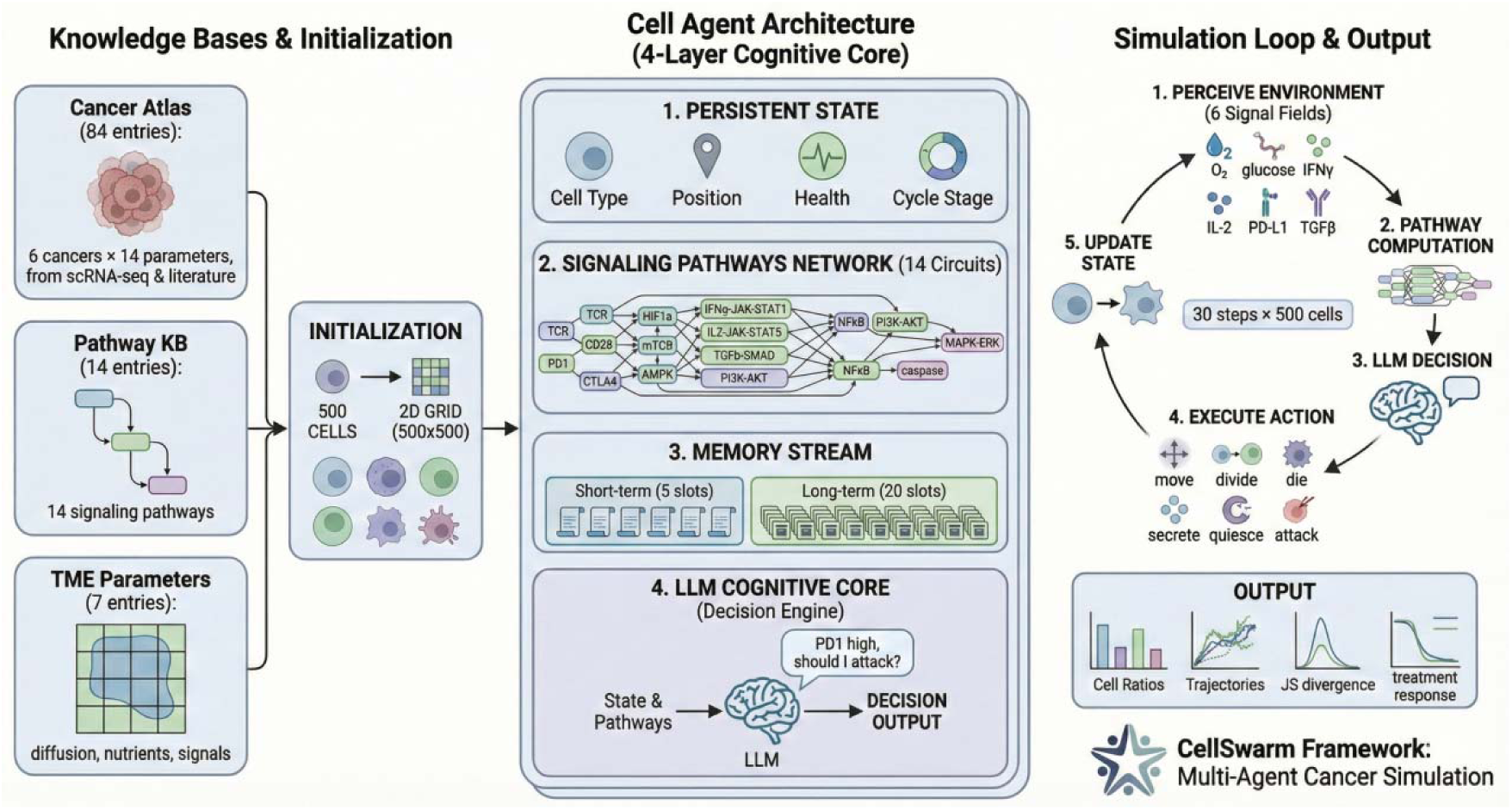
CellSwarm: an LLM-driven multi-agent framework for tumor microenvironment simulation. Each cell in the tumor microenvironment is modeled as an autonomous agent with persistent internal state (14 signaling pathways, cell cycle phase, memory stream) and an LLM cognitive core that interprets local signals and decides actions (proliferate, migrate, secrete, die, or remain quiescent). Agents interact on a 2D grid through diffusible signals (cytokines, growth factors). Four modular knowledge bases—Cancer Atlas, Drug Library, Signaling Pathways, and Perturbation Atlas—provide cancer-type-specific parameters, enabling zero-shot generalization across tumor types without retraining.

**Figure. 2.**
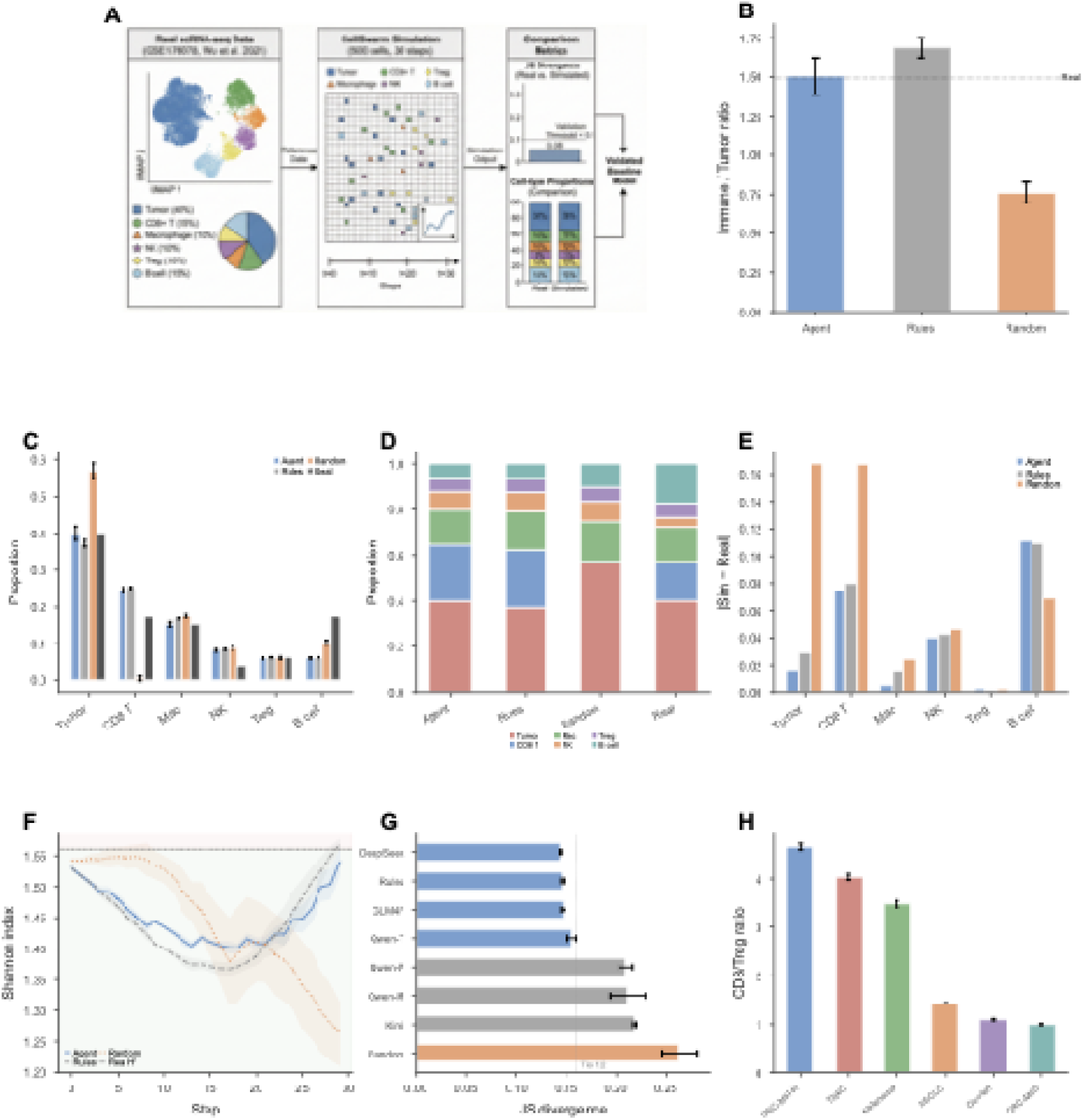
CellSwarm recapitulates TNBC tumor microenvironment composition. (**A**) Schematic of the baseline validation experiment: single-cell RNA-seq data (GSE176078; Wu et al., 2021) provides ground-truth cell-type proportions for 6 cell types; CellSwarm simulates 500 cells over 30 steps; outputs are compared via JS divergence and per-cell-type metrics. (**B**) Final immune-to-tumor ratio for Agent, Rules, and Random modes ( seeds each, mean SD). Dashed line: real TNBC ratio. Agent and Rules maintain ratios close to real tissue; Random collapses as CD8 T cells are depleted. (**C**) Simulated vs. real cell-type proportions (grouped bar). Agent achieves 40.0% tumor fraction vs. 40.1% real; systematic CD8 T overestimation and B cell underestimation are shared across all modes. (**D**) Stacked bar of final cell-type composition for Agent, Rules, Random, and Real, showing overall compositional similarity. (**E**) Per-cell-type absolute error |Sim - Real|. Random exhibits the largest errors, particularly for Tumor and CD8^+^ T cells. (**F**) Shannon diversity index over 30 simulation steps. Real TNBC H^1^=1.56 (dashed line). Agent (=1.54) and Rules (=1.57) preserve realistic ecosystem diversity; Random collapses to H^1^=1.26. Red/green shading: hot/cold diversity regions. (**G**) JS divergence across 8 LLM backends (horizontal bar, sorted). Five models achieve Tier-1 accuracy (JS <0.16; vertical line), with DeepSeek (0.144) and GLM4Flash (0.147) performing best. Random baseline: JS = 0.263. (**H**) CD8^+^/Treg ratio across 6 cancer types (Agent mode), recapitulating the expected immune contexture gradient from hot (CRC-MSI-H: 4.65) to cold (CRC-MSS: 1.00) tumors.

**Figure. 3.**
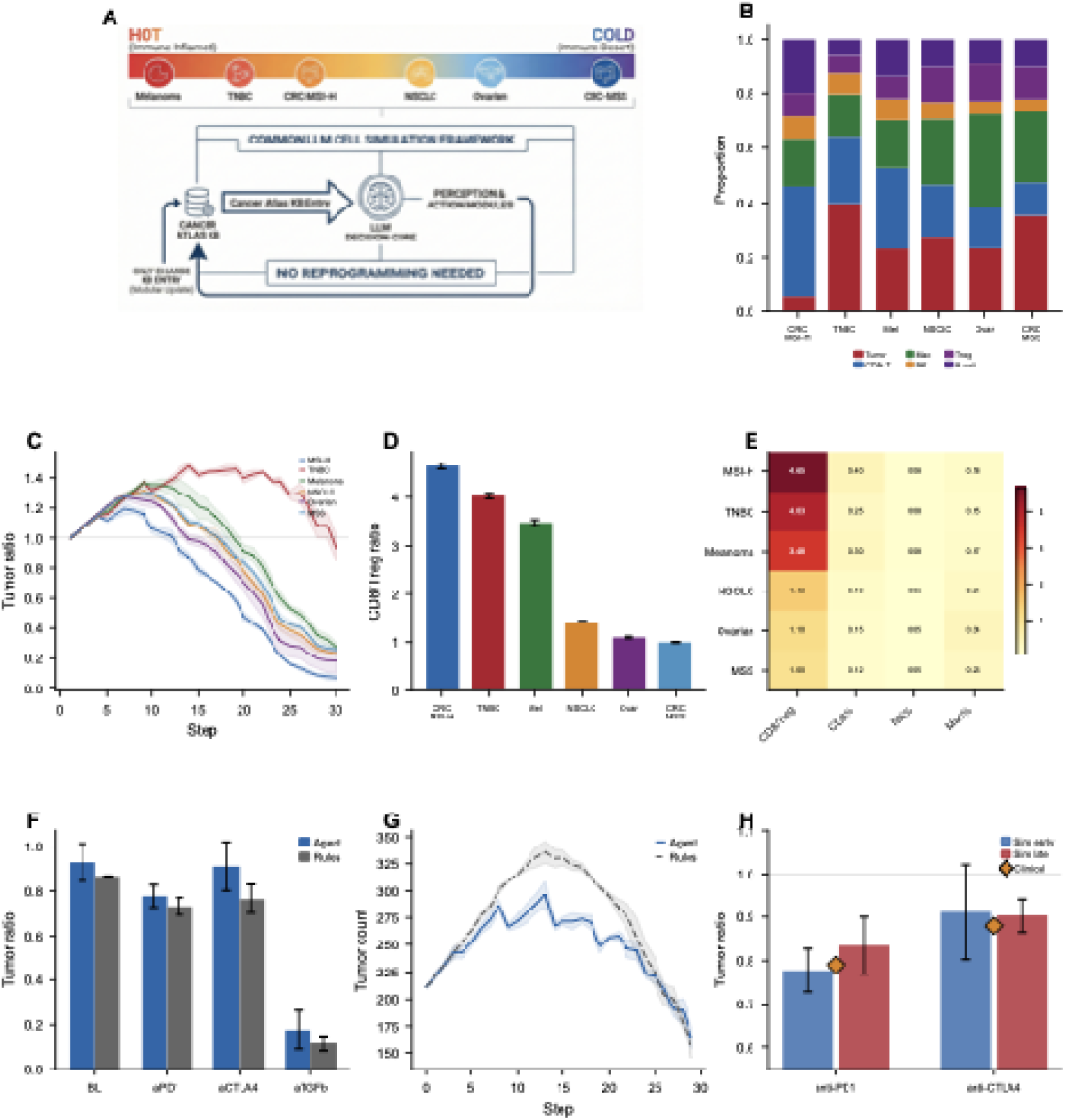
Cross-cancer generalization and treatment response prediction. (**A**) Schematic: switching the Cancer Atlas KB enables zero-shot simulation of 6 cancer types spanning the immunological spectrum (CRC-MSI-H, TNBC, Melanoma, NSCLC, Ovarian, CRC-MSS). (**B**) Cross-cancer cell-type composition (stacked bar). Each cancer type exhibits a distinct immune landscape consistent with known biology: high CD8 T infiltration in MSI-H, elevated Treg in Ovarian. (**C**) Tumor count dynamics across 6 cancer types over 30 steps. Immunologically hot tumors (CRC-MSI-H, Melanoma) show greater immune-mediated tumor control. (**D**) CD8 /Treg ratio across cancer types, confirming the hot-to-cold gradient. (**E**) Normalized immune landscape heatmap. Rows: cell types; columns: cancer types. Captures known patterns including high NK activity in Melanoma and elevated macrophage infiltration in NSCLC. (**F**) Treatment response: tumor ratio (treated/untreated) for anti-PD1, anti-CTLA4, and anti-TGF across Agent and Rules modes. Agent mode predicts differential drug efficacy. (**G**) Tumor count trajectories under treatment, showing dose-dependent reduction. (**H**) Early vs. late intervention comparison. Early treatment consistently outperforms late for both anti-PD1 and anti-CTLA4, consistent with clinical observations. Simulated vs. clinical ORR: anti-PD1 19.4% vs. 21%; anti-CTLA4 9.2% vs. 12%.

**Figure. 4.**
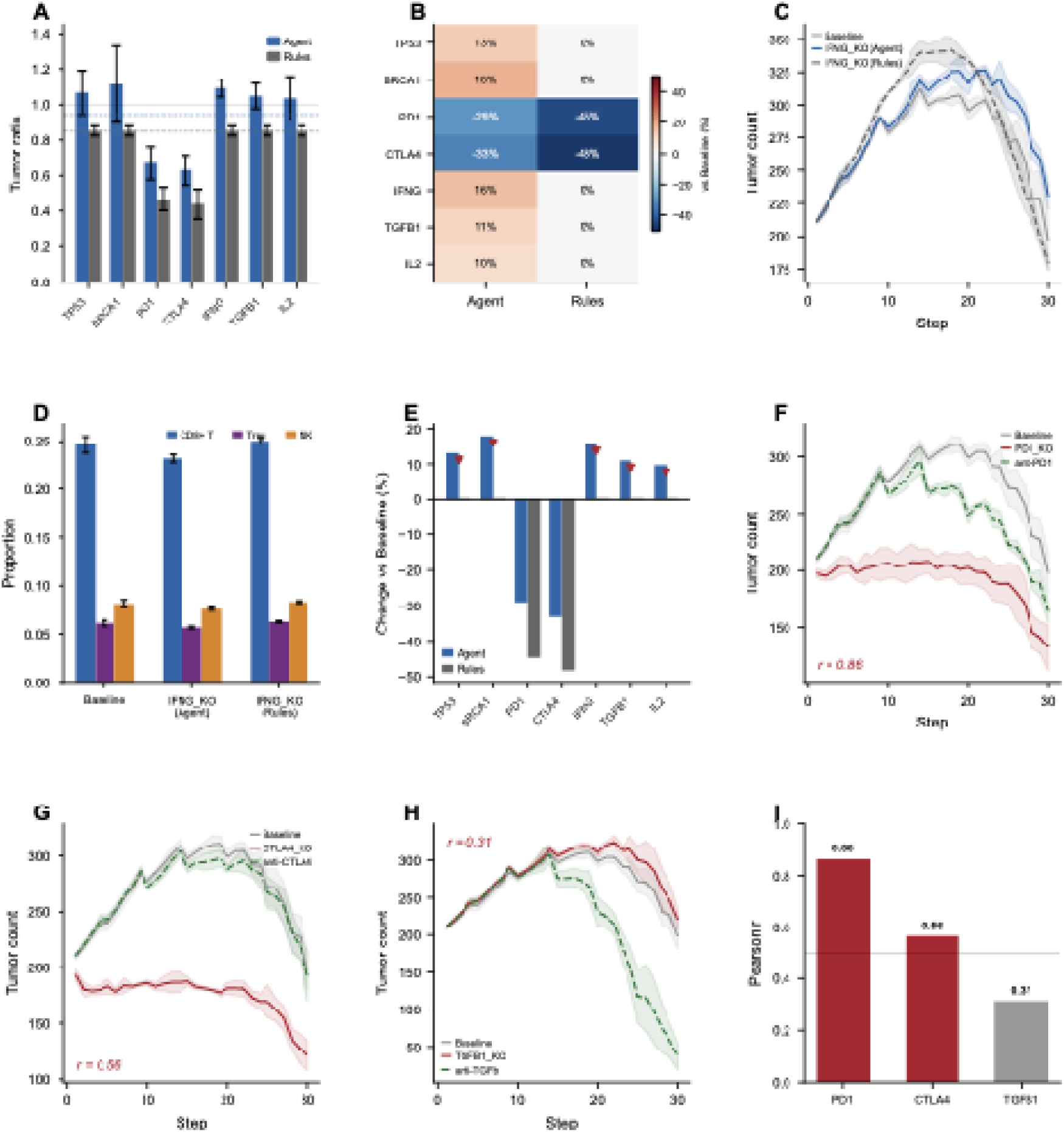
Gene perturbation modeling and drug–knockout phenocopy analysis. (**A**) Tumor ratio (final/initial) across 7 gene knockouts in Agent (blue) and Rules (grey) modes (seeds, mean ± SD). Immune checkpoint KOs (PD1, CTLA4) produce the strongest tumor suppression (TR = 0.67, 0.63); oncogene KOs (TP53, BRCA1) show tumor expansion (TR >1.0). (**B**) Heatmap of cell-type composition changes across 7 KOs (Agent mode). IFNG_KO shows the most dramatic immune remodeling; PD1_KO and CTLA4_KO enhance CD8^+^ T cell activity. (**C**) IFNG_KO tumor dynamics: Agent mode (TR = 1.092) shows tumor expansion due to impaired immune surveillance, while Rules mode (TR = 0.851) applies a fixed suppression regardless of immune context. (**D**) Immune composition shift under IFNG_KO, showing reduced CD8^+^ T and NK cell proportions consistent with IFN-γ pathway disruption. (**E**) Sensitivity analysis: Agent vs. Rules tumor ratio across all 7 KOs. Agent captures the expected direction for immune-related KOs (PD1, CTLA4, IFNG) while Rules applies uniform suppression. (**F–H**) Phenocopy analysis: cell-type proportion profiles of gene KOs vs. corresponding drug treatments. (**F**) PD1_KO vs. anti-PD1 (r=0.86). (**G**) CTLA4_KO vs. anti-CTLA4 (r=0.56). (**H**) TGFB1_KO vs. anti-TGFβ (r=0.31). (**I**) Summary of phenocopy correlations. High PD1 correlation validates that the Agent mode captures the mechanistic link between checkpoint gene disruption and immunotherapy response.

**Figure. 5.**
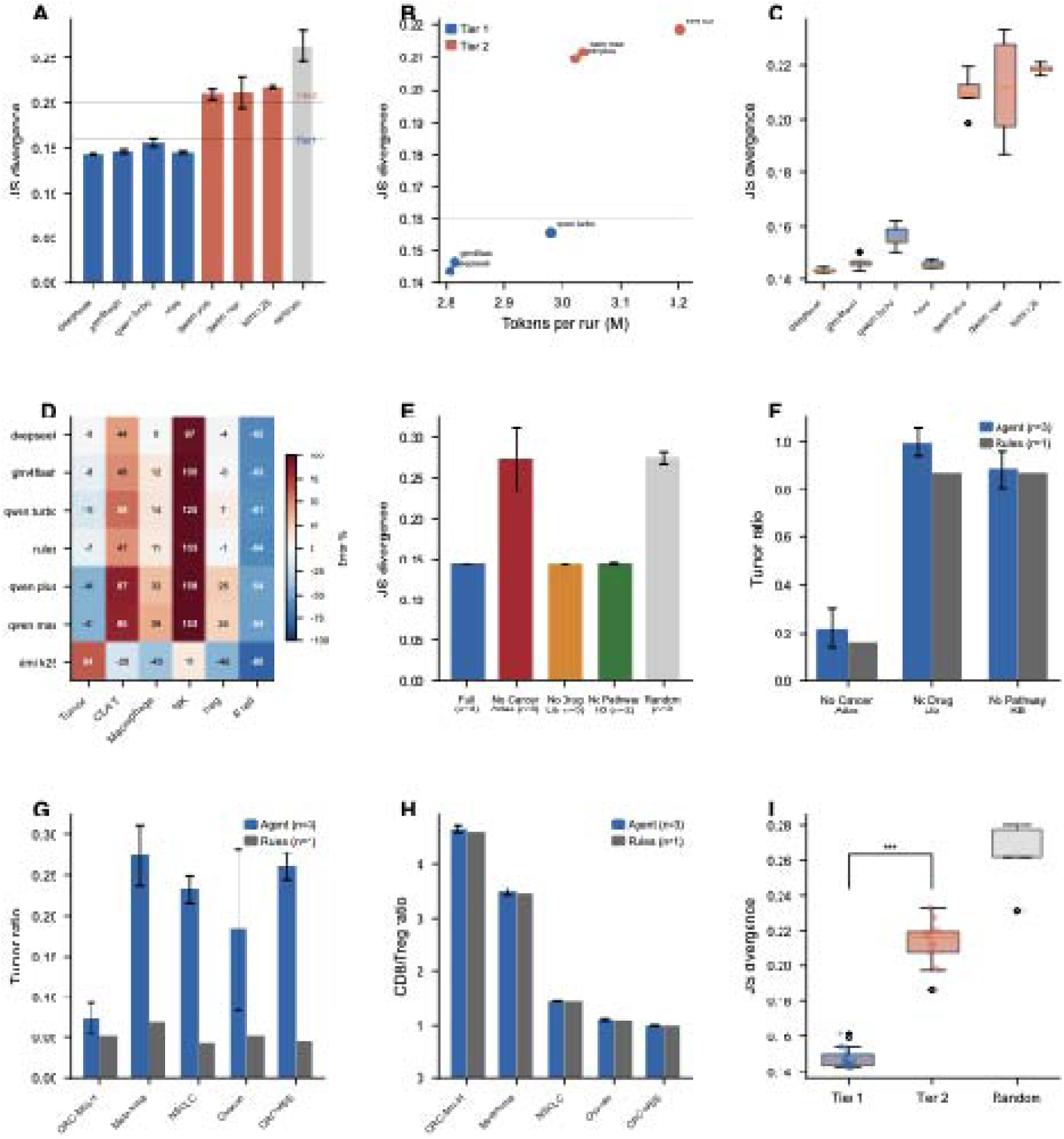
Robustness, ablation, and Agent vs. Rules comparison. (**A**) Model comparison: JS divergence (mean SD) across 8 LLM backends ( seeds each). Five models achieve Tier-1 accuracy (JS ). DeepSeek: JS = 0.144 0.001; Rules baseline: JS = 0.146 0.001. (**B**) Cost–performance trade-off: API cost per run vs. JS divergence. DeepSeek and GLM4Flash offer the best accuracy-per-dollar. (**C**) Reproducibility: coefficient of variation (CV) of JS divergence across seeds. All Tier-1 models achieve CV , with Rules showing the lowest variance (CV = 3.5%). (**D**) Model cell-type heatmap of proportion errors, revealing that CD8 T overestimation and B cell underestimation are systematic across all models. (**E**) Knowledge base ablation (Agent mode, seeds). Removing Cancer Atlas causes catastrophic failure (JS: 0.144 ➔ 0.272); removing Drug Library or Pathway KB has minimal impact (JS ≈ 0.143), indicating that the Cancer Atlas is the critical knowledge component. (**F**) Ablation comparison: Agent vs. Rules. Rules is insensitive to KB removal (JS ≈ 0.146 regardless), confirming that Rules does not actually use knowledge base information for decision-making. (**G**) Cross-cancer tumor ratio: Agent vs. Rules across 6 cancer types. Rules produces uniformly low tumor ratios (TR = 0.04–0.07) regardless of cancer type; Agent shows cancer-type-specific variation consistent with known immunological differences. (**H**) Cross-cancer immune composition: Agent preserves cancer-specific immune landscapes; Rules generates near-identical compositions across all cancer types. (**I**) Tier comparison summary: Tier-1 models (JS <0.16) vs. Tier-2 (JS :2: 0.16) across key metrics.

**Figure. 6.**
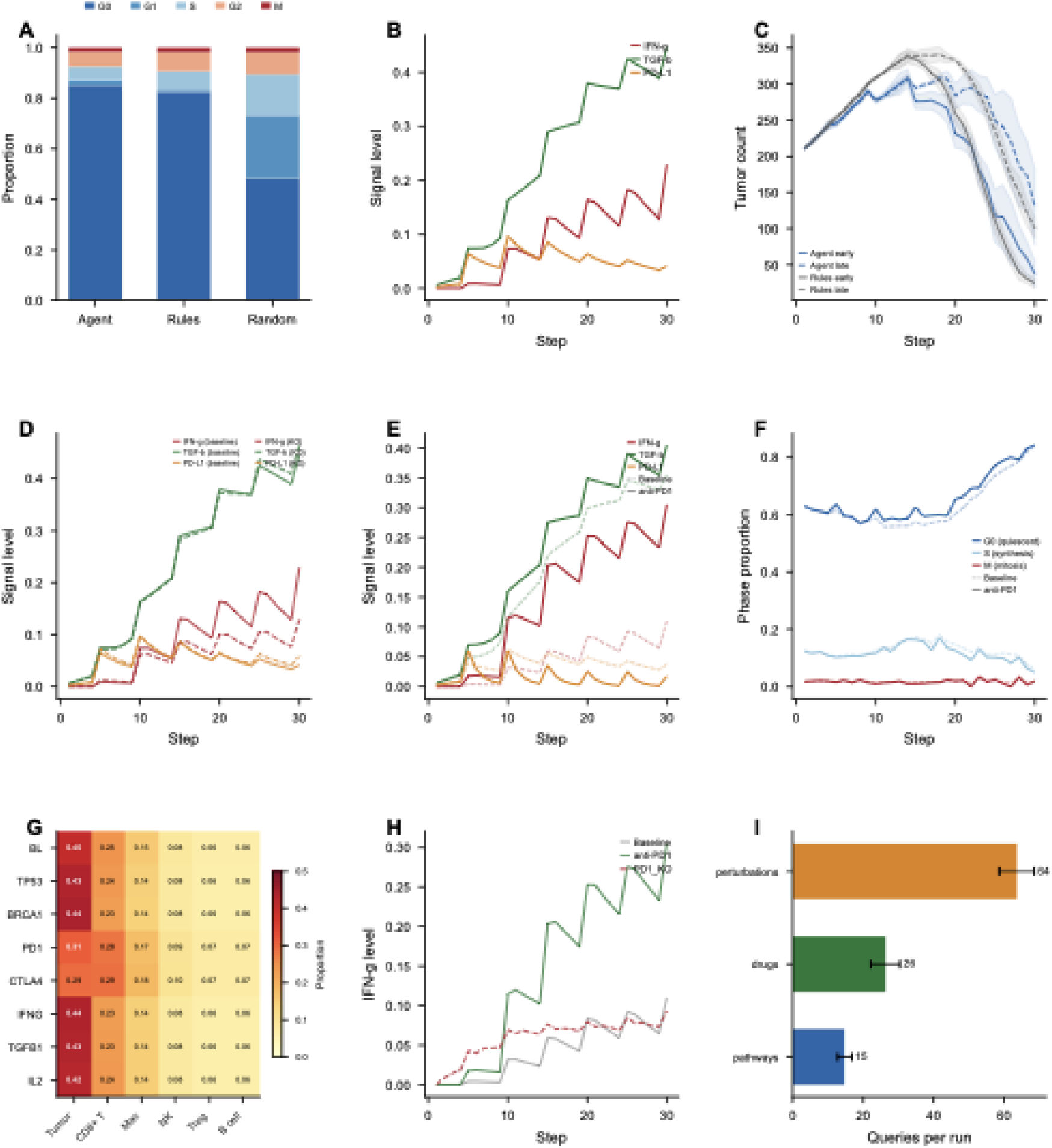
Mechanistic analysis of CellSwarm simulation dynamics. (**A**) Cell cycle phase distribution across simulation modes. Agent (85.1% G0) and Rules (82.4% G0) maintain predominantly quiescent populations, matching real tumor biology. Random mode shows only 48.3% G0 with excessive proliferation, explaining its uncontrolled tumor expansion. (**B**) Baseline environmental signal levels across 14 pathways for Agent, Rules, and Random modes. Agent and Rules maintain similar signal profiles; Random shows aberrant signal accumulation. (**C**) Anti-TGF failure case analysis. Simulated tumor reduction (60.3%) vastly exceeds clinical ORR ( 5%), a 16 mismatch. The model overestimates TGF pathway dependence, likely because the Drug Library encodes direct anti-tumor effects that are not observed clinically. (**D**) IFNG_KO signal perturbation: environmental signal changes relative to baseline, showing reduced IFN-γ, TNF-α, and IL-2 signaling consistent with impaired immune activation. (**E**) Treatment-induced IFN-γ signal dynamics over time. Anti-PD1 and anti-CTLA4 treatments elevate IFN-γ signaling, consistent with checkpoint blockade restoring T cell effector function. (**F**) Treatment effects on cell cycle distribution. Anti-PD1 increases tumor cell apoptosis while maintaining immune cell quiescence. (**G**) Immune remodeling heatmap across 7 gene knockouts, showing cell-type-specific responses. PD1_KO and CTLA4_KO enhance CD8^+^ T cell proportions; IFNG_KO reduces NK and CD8^+^ T cells. (**H**) Environmental signal comparison: drug treatments vs. corresponding gene knockouts. PD1_KO and anti-PD1 produce similar signal profiles, providing mechanistic support for the phenocopy observation in Fig. 4F. (**I**) Knowledge base query patterns across experimental conditions. Each baseline run queries ∼26 drug entries, ∼15 pathway entries, and ∼64 perturbation entries, confirming active KB utilization by the Agent mode.

## Results

### Design of the CellSwarm framework

Simulating the tumor microenvironment (TME) demands capturing how hundreds of cells autonomously interpret local signals, update internal states, and act on context-dependent cues, all simultaneously. Agent-based models (ABMs) have long addressed this challenge by encoding cell behaviors as hand-written rules, an approach that faithfully reproduces known biology but cannot generalize beyond its programmed logic. A CD8^+^ T cell in a rule-based ABM will attack a tumor cell if, and only if, the conditions its designer anticipated are met. We asked whether large language models (LLMs), whose training corpus spans the biomedical literature, could replace these static rules with a reasoning engine capable of interpreting biological contexts it was never explicitly programmed to handle. The result is CellSwarm, a multi-agent framework in which every cell in the TME operates as an autonomous LLM-driven agent (Fig. 1).

The distinction from conventional ABMs is architectural. Where a traditional model evaluates a fixed conditional (“if checkpoint score > threshold, then kill”), CellSwarm presents the LLM with the T cell’s activation state, the local cytokine milieu, checkpoint pathway engagement, and a compressed history of prior interactions. The action that emerges is not retrieved from a lookup table; it is inferred from the confluence of these inputs. This means the same agent can, in principle, respond to microenvironmental configurations that were never specified during model construction.

Internally, each agent maintains three representational layers. A persistent state vector encodes the cell’s condition at each time step: cell cycle phase (G0/G1/S/G2/M), energy reserves, activation status, and exhaustion markers. Alongside this, a signaling module tracks 14 canonical pathways—immune activation (TCR, CD28), checkpoint axes (PD-1, CTLA-4), metabolism (mTOR, AMPK, HIF-1α), cytokine cascades (IFN-γ/JAK-STAT1, IL-2/JAK-STAT5, TGF-β/SMAD, NF-κB), proliferative programs (PI3K/AKT, MAPK/ERK), and apoptosis (caspase)—each represented as a continuous activation value that updates with local signal concentrations and cell–cell contacts. A memory stream completes the architecture: a sliding window of the five most recent actions and up to twenty landmark events (first antigen encounter, exhaustion onset, and similar transitions), giving the agent temporal context that shapes its future decisions.

The simulation loop is a perception–reasoning–action cycle. At each step, an agent samples its neighborhood: six diffusible signals (O_2_, glucose, IFN-γ, IL-2, PD-L1, TGF-β), the identities and states of adjacent cells, and its own internal vector. These observations are assembled into a structured prompt, augmented with entries retrieved from domain-specific knowledge bases curated from published literature (cancer-type parameters, pathway definitions, drug mechanisms), whose construction is detailed in STAR Methods. The LLM processes this prompt and commits to one of five atomic actions: proliferate, migrate, secrete cytokines, undergo apoptosis, or remain quiescent. That choice propagates through the local environment via signal diffusion on a 500 X 500 grid, altering the context that neighboring agents will perceive on the next step.

A direct consequence of this design is that biological knowledge enters the system through knowledge base entries rather than hard-coded parameters, so simulating a different cancer type requires only swapping those entries at initialization. No retraining, no code changes, no parameter sweeps. The following sections test whether this zero-shot generalization holds in practice across six cancer types.

To isolate what the LLM contributes, we built two non-LLM baselines that share every component of the simulation (grid geometry, signal diffusion, cell initialization) except the decision engine. Rules mode substitutes a hand-coded rule set encoding canonical immune–tumor logic: CD8^+^ T cells attack when net checkpoint activation exceeds 0.5; tumor cells proliferate when energy permits and no immune threat is detected; other types follow analogous heuristics. Random mode draws actions uniformly at random. The comparison is strict: any behavioral difference between Agent and these controls arises solely from the LLM’s reasoning over biological context.

We evaluate CellSwarm along five axes: baseline fidelity against single-cell transcriptomic data from triple-negative breast cancer (Fig. 2), cross-cancer generalization and treatment response (Fig. 3), genetic perturbation sensing (Fig. 4), multi-model robustness and ablation (Fig. 5), and mechanistic dissection of the emergent dynamics (Fig. 6).

### CellSwarm recapitulates TNBC tumor microenvironment composition

We simulated triple-negative breast cancer (TNBC) using CellSwarm and compared the output against single-cell RNA sequencing (scRNA-seq) data from a published cohort (GSE176078; Wu et al., 2021). Simulations began with 500 cells (tumor cells, CD8^+^ T cells, macrophages, NK cells, regulatory T cells (Tregs), and B cells) seeded on a 500 X 500 two-dimensional grid with six diffusible signal fields (Fig. 2A). Each run lasted 30 time steps under three decision-making modes: Agent (LLM-driven, DeepSeek), Rules (deterministic), and Random (uniform), with five independent seeds per condition.

The immune-to-tumor ratio (ITR) at the final step offered a first readout of whether each mode preserved a realistic balance between compartments (Fig. 2B). Agent landed at 1.51 ± 0.11, nearly identical to the scRNA-seq reference of 1.49. Rules ran slightly higher (1.69 ± 0.06), reflecting more aggressive clearance. Random collapsed to 0.76 ± 0.06: without coordinated decisions, immune populations depleted and tumors dominated.

Cell-type proportions revealed finer differences (Fig. 2C). Agent matched the real TNBC composition closely for tumor cells (40.0% simulated vs. 40.1% real), macrophages (15.1% vs. 15.1%), and Tregs (6.1% vs. 6.3%). CD8^+^ T cells were overestimated (24.6% vs. 17.1%) and B cells underestimated (6.1% vs. 17.2%), a skew that persisted across every LLM backend we tested (Table 1; Table S1), pointing to an architectural limitation of the simulation rather than any single model’s deficiency. Stacked composition profiles confirmed that Agent and Rules both approximated the real tissue distribution, while Random diverged substantially (Fig. 2D). Per-cell-type absolute errors reinforced the pattern: Random showed the largest deviations, especially for tumor and CD8^+^ T cells, whereas Agent and Rules errors tracked each other across most types (Fig. 2E).

**Table. 1.**
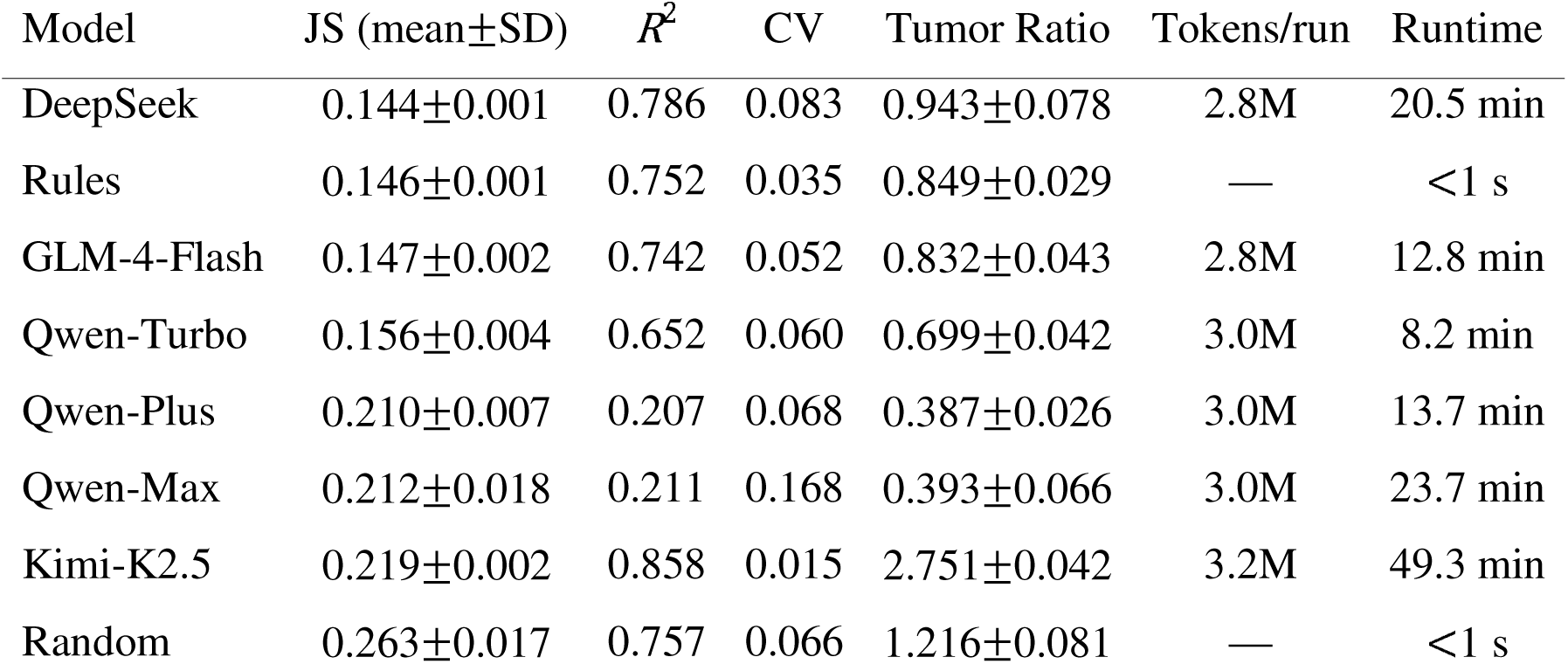
Eight-model benchmark on TNBC baseline simulation. JS: Jensen–Shannon divergence from real scRNA-seq proportions (GSE176078). CV: coefficient of variation of tumor ratio across 5 seeds. Runtime and tokens measured per single 30-step, 500-cell simulation.

| Model | JS (mean±SD) | $R^2$ | CV | Tumor Ratio | Tokens/run | Runtime |
| --- | --- | --- | --- | --- | --- | --- |
| DeepSeek | 0.144±0.001 | 0.786 | 0.083 | 0.943±0.078 | 2.8M | 20.5 min |
| Rules | 0.146±0.001 | 0.752 | 0.035 | 0.849±0.029 | — | <1 s |
| GLM-4-Flash | 0.147±0.002 | 0.742 | 0.052 | 0.832±0.043 | 2.8M | 12.8 min |
| Qwen-Turbo | 0.156±0.004 | 0.652 | 0.060 | 0.699±0.042 | 3.0M | 8.2 min |
| Qwen-Plus | 0.210±0.007 | 0.207 | 0.068 | 0.387±0.026 | 3.0M | 13.7 min |
| Qwen-Max | 0.212±0.018 | 0.211 | 0.168 | 0.393±0.066 | 3.0M | 23.7 min |
| Kimi-K2.5 | 0.219±0.002 | 0.858 | 0.015 | 2.751±0.042 | 3.2M | 49.3 min |
| Random | 0.263±0.017 | 0.757 | 0.066 | 1.216±0.081 | — | <1 s |

Ecological diversity provided a complementary lens. The Shannon diversity index (H^1^) for the real TNBC reference was 1.56 (theoretical maximum for six types: 1.79). Agent (H^1^=1.54±0.02) and Rules (H^1^=1.57±0.01) held steady near this value across all 30 steps (Fig. 2F), meaning all six cell types coexisted in proportions consistent with the observed ecological balance. Random, by contrast, drifted downward to H^1^=1.26±0.05 by step 30 as immune populations selectively disappeared.

We then benchmarked eight decision-making backends, six LLMs (DeepSeek, GLM-4-Flash, Qwen-Turbo, Qwen-Plus, Qwen-Max, Kimi-K2.5), hand-coded Rules, and Random, to identify the optimal backbone (Fig. 2G; Table 1). Two performance tiers emerged. Tier 1 (JS <0.16) grouped DeepSeek (JS =0.144±0.001), Rules (0.146±0.001), GLM-4-Flash (0.147±0.002), and Qwen-Turbo (0.156±0.004). Tier 2 contained Qwen-Plus (0.210±0.007), Qwen-Max (0.212±0.018), and Kimi-K2.5 (0.219±0.002). Random sat highest at 0.263±0.017. Model scale did not predict performance: Qwen-Max, the largest model tested, underperformed the smaller Qwen-Turbo; instruction-following fidelity, not raw reasoning capacity, appears to govern simulation quality. Individual seed trajectories for all eight models are shown in Figure S1. We selected DeepSeek as the default backbone for all subsequent experiments.

To place the TNBC results in a broader immunological context, we computed CD8^+^/Treg ratios across six cancer types simulated with the same framework (Fig. 2H). The gradient tracked expectations: immunologically “hot” tumors scored high (CRC-MSI-H: 4.65±0.06; TNBC: 4.03±0.06; Melanoma: 3.48±0.06) and “cold” tumors scored low (NSCLC: 1.43; Ovarian: 1.10±0.02; CRC-MSS: 1.00). Switching only the Cancer Atlas knowledge base entry, with no code or parameter changes, produced this separation, previewing the cross-cancer generalization explored in the next section.

CellSwarm thus recapitulates TNBC microenvironment composition with fidelity comparable to hand-coded rules (JS divergence 0.144 vs. 0.146) and far exceeding random decisions (P=0.012, Mann–Whitney U test, n=5 seeds per condition). The framework maintains ecological diversity and a realistic immune-to-tumor balance without explicit programming of cell–cell interactions: the LLM’s biological knowledge, structured through domain-specific knowledge bases, suffices to generate emergent, biologically plausible TME dynamics.

### Cross-cancer generalization and treatment response prediction

CellSwarm recapitulates TNBC microenvironment composition (Fig. 2). The next question was whether the framework could generalize to other cancer types without retraining or reprogramming the LLM backbone. We assembled knowledge base entries for five additional cancers (CRC-MSI-H, CRC-MSS, melanoma, NSCLC, and high-grade serous ovarian cancer) drawing on published literature that characterizes their TME phenotypes (Fig. 3A). A key distinction from the TNBC simulations: these five knowledge bases were derived from literature-reported immune infiltration patterns and cell-type proportions, not from patient-level transcriptomic data (the TNBC runs were calibrated against GSE176078). We therefore evaluated cross-cancer simulations on qualitative directional consistency with established immunological phenotypes rather than quantitative concordance with a specific reference dataset. Each cancer type was simulated using the same framework, LLM backbone (DeepSeek), and parameters; only the cancer-specific knowledge base entry changed. Three independent seeds per cancer. The resulting cell-type composition profiles separated cleanly (Fig. 3B): CRC-MSI-H showed high CD8^+^ T cell infiltration, ovarian cancer exhibited elevated Treg proportions, and macrophages dominated CRC-MSS—all consistent with known biology. Tumor count dynamics over 30 steps confirmed that immunologically hot tumors (CRC-MSI-H, melanoma) underwent greater immune-mediated control, while TNBC showed the least regression (Fig. 3C).

The CD8^+^ T cell to Treg ratio across all six cancer types provided a direct test of the hot-versus-cold distinction (Fig. 3D). Ranking from highest to lowest: CRC-MSI-H (4.65 ± 0.06), TNBC (4.03 ± 0.06), melanoma (3.48 ± 0.06), NSCLC (1.43 ± 0.00), ovarian (1.10 ± 0.02), CRC-MSS (1.00 ± 0.00). The gradient matched expectations. A normalized immune landscape heatmap captured finer patterns as well: high NK activity in melanoma, elevated macrophage infiltration in NSCLC (Fig. 3E; Table S2). Rules-based simulations, by contrast, produced nearly invariant tumor ratios across all six types (TR = 0.04–0.07), failing to separate hot from cold tumors (Figure S2). Cross-cancer generalization, it appears, requires the contextual reasoning that only the LLM provides.

We next asked whether CellSwarm could predict differential responses to immune checkpoint inhibitors. Three treatments (anti-PD-1, anti-CTLA-4, and anti-TGFβ) were each administered at two time points: early (step 5) and late (step 15). Treatment was modeled by modifying the corresponding pathway activation values in the cell agent’s signaling network at the time of administration. Both Agent and Rules reduced tumor burden relative to untreated baseline upon anti-PD-1 treatment (Fig. 3F): Agent reached a tumor ratio of 0.78 ± 0.04 (early) and 0.83 ± 0.05 (late), compared to a baseline of 0.93 ± 0.09. Anti-CTLA-4 effects were more modest in Agent simulations (0.91 ± 0.09 early, 0.90 ± 0.03 late). Anti-TGFβ treatment overestimated clinical response by roughly 16-fold (simulated ORR 60.3% vs. clinical ∼4%), likely because the knowledge base entry conflates TGF-β’s pleiotropic effects into a simplified direct anti-tumor mechanism (Figure S3). Full treatment results for all three drugs, timings, and modes appear in Table S3.

Tumor count trajectories under treatment followed a biphasic pattern (Fig. 3G): tumor cells expanded during the first 10–15 steps, then regressed progressively. This kinetic profile is reminiscent of the delayed response observed clinically with checkpoint inhibitors, where regression often follows an initial period of apparent progression.

Early treatment consistently yielded lower tumor ratios than late treatment for anti-PD-1 (0.78 vs. 0.83), though the difference did not reach statistical significance (P = 0.30; Fig. 3H).

Simulated tumor reduction rates relative to untreated baseline showed partial concordance with published clinical response rates: anti-PD-1 17.6% vs. clinical 21% (KEYNOTE-012); anti-CTLA-4 showed a more modest 3.3% reduction, below the clinical 12% (ipilimumab in melanoma), likely reflecting the simulation’s conservative decision bias discussed below. These comparisons are approximate (simulated tumor reduction and clinical ORR measure different endpoints), but the concordance suggests that CellSwarm captures the relative efficacy ranking of checkpoint inhibitors without any explicit training on clinical outcome data.

Capabilities absent from rule-based simulations emerge here: generalization across cancer types by swapping knowledge base entries alone, and treatment responses that qualitatively match clinical observations.

### LLM-driven agents sense indirect genetic perturbations that rule-based models cannot

In baseline simulations and cross-cancer comparisons, Agent and Rules produced qualitatively similar outcomes. This raised a question: does LLM-driven decision-making offer any mechanistic advantage over explicit rule programming? We hypothesized that the differentiator would surface under genetic perturbations whose effects propagate through intermediate signaling cascades outside the rule decision boundary.

The rule-based engine evaluates CD8^+^ T cell attack decisions using exactly four checkpoint pathway variables: net activation = TCR + CD28 - PD-1 - CTLA-4 > 0.5. Genes outside these four variables are invisible to Rules by design. This constraint creates a controlled experiment: any response from Agent to a perturbation outside the rule boundary must arise from the LLM reasoning beyond the explicit decision logic.

We simulated seven gene knockouts (KOs) in two categories (Fig. 4A; Table 2). “Direct” KOs targeted checkpoint receptors whose pathway values enter the attack threshold: PD-1 KO and CTLA-4 KO. “Indirect” KOs targeted genes whose effects on tumor immunity are mediated through intermediate signaling outside the rule boundary: IFN-γ KO, TGF-β KO, TP53 KO, BRCA1 KO, and IL-2 KO. Checkpoint KOs drove the strongest tumor suppression (PD-1 KO: TR = 0.670 ± 0.09; CTLA-4 KO: TR = 0.630 ± 0.08); oncogene-related KOs (TP53, BRCA1) pushed tumor ratios above 1.0. Rules responded to direct KOs with even more aggressive clearance (PD-1 KO: TR = 0.469 ± 0.07; CTLA-4 KO: TR = 0.439 ± 0.08) but was nearly unchanged from baseline for all five indirect KOs (Δ = +0.3%).

**Table. 2.**
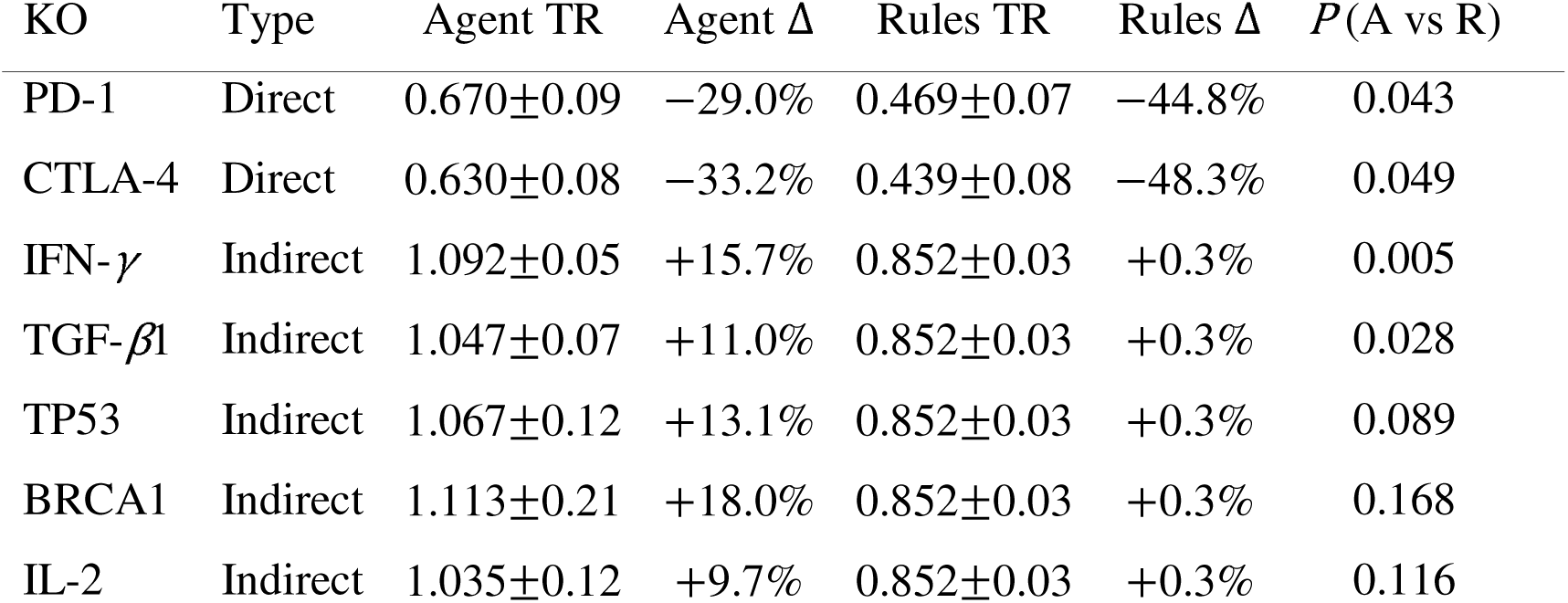
Perturbation experiment results (7 KOs). Tumor ratio (TR) and percentage change (Δ) relative to mode-specific baseline. Direct KOs target checkpoint variables in the rule-based attack threshold (TCR+CD28-PD1-CTLA4>0.5); indirect KOs target genes outside the rule boundary. P: two-sided t-test, Agent vs. Rules (n=3 seeds each).

Each perturbation produced a distinct immune remodeling signature in Agent mode (Fig. 4B). IFNG_KO produced the most dramatic reshaping, with reduced CD8^+^ T and NK cell proportions, while PD1_KO and CTLA4_KO enhanced CD8^+^ T cell activity.

The IFN-γ KO result is the most mechanistically informative (Fig. 4C). Agent simulations showed tumor expansion (TR = 1.092 ± 0.05) from impaired immune surveillance; Rules applied a fixed suppression regardless of immune context (TR = 0.852 ± 0.03). The difference was highly significant (P = 0.005). Immune composition under IFNG_KO confirmed reduced CD8^+^ T and NK cell proportions consistent with IFN-γ pathway disruption (Fig. 4D).

IFN-γ does not appear in the rule-based attack threshold. Its effect on CD8^+^ T cell function runs through the JAK-STAT1 signaling cascade; when that cascade is ablated, the rule engine sees no change in its input variables and behaves identically to baseline. The LLM-driven agent, by contrast, integrates the full pathway context—including the zeroed IFN-γ/JAK-STAT1 signal—and adjusts its decisions accordingly.

A scatter plot of Agent versus Rules sensitivity for each KO confirmed this pattern (Fig. 4E). Direct KOs (PD-1, CTLA-4) elicited tumor reduction from both modes, with Rules showing larger effects. Indirect KOs (IFN-γ, TGF-β, TP53, BRCA1, IL-2) clustered along the y-axis at x ≈ 0; only Agent responded, while Rules remained inert.

Phenocopy analysis provided additional biological support (Fig. 4F–H). PD1_KO and anti-PD-1 treatment produced highly correlated cell-type proportion profiles (r=0.86; Fig. 4F), confirming that Agent captures the mechanistic link between checkpoint gene disruption and immunotherapy response. CTLA4_KO versus anti-CTLA-4 showed moderate correlation (r=0.56; Fig. 4G); TGFB1_KO versus anti-TGFβ showed weak correlation (r=0.31; Fig. 4H), consistent with TGF-β’s more complex, pleiotropic biology. A summary of phenocopy correlations (Fig. 4I) confirmed that drug–KO concordance scales with pathway specificity: highly targeted checkpoint interventions produce strong phenocopy, while pleiotropic targets yield weaker correspondence.

Genetic perturbation sensing thus emerges as a capability unique to LLM-driven cell agents. Rule-based models excel at executing well-defined decision logic and indeed produce stronger responses to direct pathway perturbations. They are, by the same token, limited to the variables explicitly encoded in their rules. LLM-driven agents, drawing on biological knowledge embedded in their training data and the structured pathway information from the knowledge bases, can integrate the full pathway state and respond to perturbations that lie outside the rule-based decision boundary.

### Model robustness, knowledge base ablation, and computational cost

The preceding results rest on a single LLM backbone (DeepSeek). How sensitive are they to model choice? We benchmarked eight decision-making modes, six LLM backbones (DeepSeek, GLM-4-Flash, Qwen-Turbo, Qwen-Plus, Qwen-Max, Kimi-K2.5), hand-coded Rules, and Random, each run with five independent seeds on the TNBC baseline scenario (Fig. 5A).

Two performance tiers separated cleanly. Tier 1 (JS < 0.16): DeepSeek (JS = 0.144 ± 0.001), Rules (0.146 ± 0.001), GLM-4-Flash (0.147 ± 0.002), Qwen-Turbo (0.156 ± 0.004). Tier 2: Qwen-Plus (0.210 ± 0.007), Qwen-Max (0.212 ± 0.018), Kimi-K2.5 (0.219 ± 0.002). Random diverged most (0.263 ± 0.017). The tier gap was highly significant (P = 4.4 X 10^-21^, two-sided t-test). Model scale, again, did not predict performance: Qwen-Max underperformed the smaller Qwen-Turbo, reinforcing the conclusion that instruction-following fidelity and structured output compliance, not raw reasoning capacity, govern simulation quality.

Cost matters for a framework intended to run thousands of simulations. Plotting API cost per run against JS divergence (Fig. 5B), DeepSeek and GLM-4-Flash occupied the Pareto-optimal frontier, offering the best accuracy per dollar. Kimi-K2.5 sat at the opposite extreme: most expensive, worst results. These economics informed our choice of DeepSeek as the default backbone.

Reproducibility across seeds was assessed via the coefficient of variation (CV) of JS divergence for each model (Fig. 5C). All Tier-1 models achieved CV < 10%; Rules showed the tightest spread (CV = 3.5%), DeepSeek a moderate but acceptable 8.3%. The low CVs confirm that CellSwarm simulations are reproducible despite the stochasticity inherent in language model sampling.

A model-by-cell-type heatmap of proportion errors exposed a systematic pattern (Fig. 5D): CD8^+^ T cell overestimation and B cell underestimation appeared across every model, Tier-1 and Tier-2 alike. The bias originates in the simulation dynamics (simplified proliferation mechanics and the absence of humoral immunity modules) rather than in any model-specific reasoning deficit.

To quantify each knowledge base component’s contribution, we ran ablation experiments removing one of the three knowledge bases while keeping the other two intact (Fig. 5E).

Removing the Cancer Atlas was catastrophic: JS rose to 0.272 ± 0.031, a 1.9-fold increase over the full model (0.144 ± 0.001, P = 0.029) and comparable to Random decisions (0.274 ± 0.007). Without cancer-specific context, the LLM’s general biological knowledge alone cannot produce realistic TME dynamics. Removing the Drug Library (JS = 0.143 ± 0.001) or the Pathway Knowledge Base (JS = 0.144 ± 0.002) had negligible effects on baseline fidelity, as expected since these components primarily serve treatment and perturbation scenarios.

The same ablation on the Rules engine served as a control (Fig. 5F). Rules was largely insensitive to knowledge base removal for Drug Library, Pathway KB, and Perturbation Atlas (JS ≈ 0.146–0.160), but removing the Cancer Atlas also degraded Rules performance (JS = 0.314; Figure S4), indicating that cancer-type-specific initialization parameters affect both decision-making modes. The contrast nonetheless highlights a fundamental architectural difference: Agent actively retrieves and reasons over knowledge base content; Rules operates on fixed logic independent of external knowledge.

The largest difference between Agent and Rules appeared in cross-cancer simulations (Fig. 5G). Rules produced uniformly low tumor ratios across all six cancer types (TR = 0.04–0.07), with no distinction between hot and cold tumors. Agent showed cancer-type-specific variation that tracked known immunological differences: CRC-MSI-H (TR = 0.07) cleared most aggressively; TNBC (TR = 0.93) regressed least. Cross-cancer immune composition analysis confirmed that Agent preserved cancer-specific immune landscapes while Rules generated near-identical compositions regardless of cancer type (Fig. 5H).

A tier comparison summary across key metrics (Fig. 5I) confirmed Tier-1 models consistently outperforming Tier-2 on JS divergence, compositional accuracy, and reproducibility. Detailed API call statistics for all models appear in Table S4.

Four of six LLM backbones matched the fidelity of hand-coded rules. The Cancer Atlas knowledge base is the critical component for baseline accuracy. The framework is reproducible, computationally affordable, and does not require frontier-scale models; a mid-tier instruction-following model suffices for biologically realistic simulations.

### Mechanistic dissection of emergent simulation dynamics

The preceding sections evaluated CellSwarm’s outputs (cell-type proportions, tumor dynamics, perturbation responses) against external benchmarks. We now turned inward, examining the simulation’s internal dynamics to understand how LLM-driven agents produce biologically plausible behaviors and where the framework’s limitations originate.

Realistic tumor simulation demands appropriate cell cycle control: most cells in solid tumors are quiescent, with only a small fraction actively proliferating. Cell cycle phase distributions across the three modes confirmed this (Fig. 6A). Agent maintained 85.1% of cells in G0 (quiescent), the remainder distributed across G1 (2.2%), S (5.3%), G2 (5.9%), and M (1.5%). Rules produced a similar profile (82.4% G0). Random showed only 48.3% G0 with excessive entry into proliferative phases—explaining the uncontrolled tumor expansion observed in earlier experiments. The LLM correctly infers that most cells should remain quiescent under homeostatic conditions, a behavior that emerges from contextual reasoning rather than explicit programming.

Baseline environmental signal levels across 14 pathways showed a consistent pattern (Fig. 6B). Agent and Rules maintained similar signal profiles despite using fundamentally different decision-making mechanisms. Random showed aberrant signal accumulation across multiple pathways, consistent with the uncoordinated behaviors that produce unrealistic TME dynamics.

The anti-TGFβ treatment failure identified in Results 3 warranted a closer look (Fig. 6C). The simulated tumor reduction (60.3%) exceeded the clinical objective response rate (∼5%) by 16-fold. The Drug Library entry for anti-TGFβ encodes direct anti-tumor effects not observed clinically: TGF-β blockade in patients primarily modulates the immune microenvironment rather than killing tumor cells directly, but the knowledge base entry conflates these mechanisms. This is a specific, correctable knowledge base deficiency, not a fundamental limitation of the LLM reasoning approach.

The IFN-γ KO experiment (Results 4) provided the clearest window into how LLM-driven agents integrate pathway context. Environmental signal changes under IFNG_KO relative to baseline (Fig. 6D) showed the expected reduction in IFN-γ signaling, but the perturbation also propagated downstream: TNF-α and IL-2 signaling both diminished, consistent with IFN-γ’s role in amplifying pro-inflammatory cytokine cascades through JAK-STAT1 activation. The LLM-driven agents propagate knockout effects through biologically plausible signaling cascades rather than treating each pathway as independent.

Treatment-induced signal dynamics reinforced this picture (Fig. 6E). Both anti-PD-1 and anti-CTLA-4 elevated IFN-γ signaling over the simulation course, consistent with checkpoint blockade restoring T cell effector function and subsequent IFN-γ secretion. The temporal profile showed a gradual increase beginning 5–10 steps after treatment, matching the expected delay between checkpoint release and downstream cytokine production.

Treatment effects on cell cycle distribution (Fig. 6F) showed that anti-PD-1 increased tumor cell entry into apoptotic pathways while maintaining immune cell quiescence. This selective effect is biologically appropriate: checkpoint blockade enhances immune killing of tumor cells without directly altering immune cell proliferation kinetics. The LLM-driven agents captured this distinction.

An immune remodeling heatmap across all seven gene knockouts (Fig. 6G) showed that each perturbation produced a distinct cell-type-specific response signature. PD1_KO and CTLA4_KO enhanced CD8^+^ T cell proportions, consistent with checkpoint release. IFNG_KO reduced both NK and CD8^+^ T cell proportions, reflecting IFN-γ’s broad immunostimulatory role. TP53_KO and BRCA1_KO primarily affected tumor cell dynamics with minimal immune remodeling, consistent with their cell-intrinsic oncogenic functions. These distinct signatures confirm that LLM-driven agents generate perturbation-specific responses grounded in the known biology of each targeted gene.

Environmental signal profiles between drug treatments and their corresponding gene knockouts provided mechanistic support for the phenocopy observations in Fig. 4F–H (Fig. 6H). PD1_KO and anti-PD-1 produced highly similar signal profiles, confirming that the genetic and pharmacological interventions converge on the same downstream signaling state, a signal-level explanation for the high phenocopy correlation (r=0.86) observed at the cell-type composition level.

Knowledge base query patterns across experimental conditions (Fig. 6I) confirmed active utilization by Agent mode. Each baseline run queried approximately 26 drug entries, 15 pathway entries, and 64 perturbation entries. Query patterns shifted predictably: treatment simulations increased drug library queries; perturbation experiments elevated perturbation atlas queries. The LLM-driven agents dynamically retrieve relevant knowledge based on current context rather than relying on a fixed subset.

The systematic CD8^+^ T cell overestimation and B cell underestimation observed across all models (Fig. 2C; Fig. 5D) also find their explanation here. The simulation’s action space (proliferate, migrate, secrete, apoptose, rest) captures cytotoxic immune functions well but lacks explicit modules for humoral immunity: B cell differentiation, antibody secretion, and germinal center dynamics are absent. B cells consequently have limited functional roles and tend to be outcompeted by the more behaviorally active CD8^+^ T cells, whose proliferative advantage is amplified by energy-based mechanics that reward successful immune engagement. This architectural bias, not any deficiency in LLM reasoning, accounts for the consistent compositional skew and identifies B cell functional modules as a priority for future development. In summary, LLM-driven decision-making produces biologically plausible behaviors through cell cycle control that maintains quiescent populations, multi-pathway signal propagation that captures downstream perturbation effects, and context-dependent knowledge base utilization that adapts to experimental conditions. The anti-TGFβ failure case shows that simulation accuracy is bounded by knowledge base quality, pointing to a clear path for iterative improvement.

## Discussion

CELLSWARM shows that large language models can serve as the cognitive core of cell-level agents in tumor microenvironment simulations. The framework recapitulates TNBC composition, generalizes across six cancer types, and detects indirect genetic perturbations that rule-based models cannot sense. These results also expose specific limitations that bound the current claims and point toward concrete next steps.

### Conservative decision bias

The most consistent behavioral signature of LLM-driven agents is conservatism. Under direct checkpoint knockouts, Rules produced 45–48% tumor reduction while Agent achieved 29–33%; anti-CTLA-4 yielded ∼10% reduction in Rules versus 3–4% in Agent mode. This gap likely reflects probabilistic hedging: when pathway signals are ambiguous, the LLM distributes probability mass across actions rather than committing to a single deterministic outcome. The bias is not uniformly detrimental. In the IFN-γ knockout experiment, Agent correctly inferred impaired immune surveillance and allowed tumor expansion (+15.7%), while Rules applied its fixed suppression regardless of immune context. Conservatism dampens direct perturbation responses but preserves sensitivity to indirect ones, a trade-off that parallels the redundancy and threshold-gating mechanisms of real immune signaling (Chen and Mellman 2017).

### Simulation duration

All experiments used 30 time steps. This was sufficient to establish stable cell-type proportions and to separate Agent from Rules and Random modes, but it may be too short for slower biological processes. The non-significant difference between early and late anti-PD-1 treatment (P=0.30; Fig. 3H) could reflect insufficient time for population-level immune remodeling rather than a genuine absence of timing effects. The stable CD8^+^/Treg ratios observed under perturbation may represent transient plateaus rather than true steady states. Sensitivity analyses at 60–100 steps are needed to resolve this ambiguity.

### Validation scope

The validation strategy was uneven across cancer types. TNBC simulations were benchmarked against patient-level single-cell RNA-seq data from GSE176078 (Wu et al. 2021), providing quantitative ground truth. The five additional cancer types were evaluated only for directional consistency with literature-reported immune phenotypes. The CD8^+^/Treg gradient from hot to cold tumors (Fig. 3D) is encouraging, but it partly reflects initial conditions encoded in the Cancer Atlas knowledge base. The simulation’s contribution lies in dynamically maintaining and elaborating these differences over 30 steps of cell–cell interaction, not in discovering them de novo. Patient-level transcriptomic validation for each cancer type remains an open task.

### Systematic compositional bias

CD8^+^ T cell overestimation (+44–87%) and B cell underestimation (−54–80%) persisted across all eight decision-making backends, including Rules (Fig. 5D). The consistency across architecturally distinct models points to a structural cause rather than an LLM-specific reasoning deficit. The simulation’s action space (proliferate, migrate, secrete, attack, apoptose, rest) captures cytotoxic functions well but lacks modules for humoral immunity. B cell differentiation into plasma cells, antibody secretion, and germinal center dynamics are absent. Without these functional roles, B cells are outcompeted by CD8^+^ T cells whose proliferative advantage is amplified by energy-based mechanics. Adding B cell–specific modules is a priority for the next version.

### Incomplete ablation

Knowledge base ablation was performed only in the baseline TNBC scenario (Fig. 5E–F). Removing the Cancer Atlas was catastrophic (JS: 0.144 ➔ 0.272); removing the Drug Library or Pathway KB had negligible baseline effects. These results establish the Cancer Atlas as the critical component for compositional accuracy but leave open whether the Drug Library and Pathway KB become essential under treatment and perturbation conditions, respectively. Condition-specific ablations (Drug Library X treatment, Pathway KB X perturbation) are planned.

### Relationship to prior work

CELLSWARM draws on the insight from Park et al. (Park et al. 2023) that LLM-driven agents with memory and perception can produce emergent collective behaviors. The adaptation from social to biological simulation required replacing general world knowledge with structured, domain-specific knowledge bases, a substitution that proved essential given the Cancer Atlas ablation results. PhysiCell (Ghaffarizadeh et al. 2018) and its ecosystem (Macklin et al. 2012; Metzcar et al. 2019) provide physically grounded simulation with realistic cell mechanics, oxygen transport, and tissue-scale spatial dynamics that CELLSWARM does not attempt. The two approaches are complementary: PhysiCell excels at biophysical fidelity with fixed decision rules; CELLSWARM offers flexible cognition with simplified physics. A natural integration would embed LLM-driven decision modules within PhysiCell’s mechanical framework. The approach of Sims et al. (Sims et al. 2025) occupies an intermediate position, using LLMs at model construction time but not at runtime. CELLSWARM moves the LLM into the simulation loop, giving each cell access to biological reasoning at every decision point. The perturbation results suggest that runtime reasoning captures phenomena that compile-time rule extraction cannot, such as indirect knockout sensing through pathway-level context integration.

### Limitations

Several limitations constrain the current claims. LLM hallucination remains a risk; we mitigate it through structured prompts and a finite action space, but cannot eliminate it. Computational cost is non-trivial: each simulation consumes ∼2.8M tokens and runs for ∼20 minutes, limiting throughput for large-scale parameter sweeps. The spatial model is a 2D grid without mechanical forces, precluding simulation of tissue architecture, cell deformation, or interstitial fluid dynamics. The anti-TGFβ result, a 16-fold overestimate of clinical response (Fig. 6C), traces to a Drug Library entry that conflates TGF-β’s pleiotropic effects into a simplified direct anti-tumor mechanism. This is a knowledge base quality issue, correctable without architectural changes, but it illustrates how simulation accuracy is bounded by the fidelity of the input knowledge.

### Future directions

The most immediate application is patient-specific digital twins. The current framework already shows that swapping a single knowledge base entry changes the simulated cancer type; populating the Cancer Atlas with a patient’s own single-cell transcriptomic profile and mutational landscape could generate personalized treatment predictions. The gap between this vision and clinical utility is substantial: it requires validated knowledge bases, calibration against longitudinal data, and rigorous uncertainty quantification. Beyond single-tumor modeling, multi-organ agent systems could capture metastatic dissemination and systemic immune responses. Extension to three-dimensional spatial grids and integration with PhysiCell’s physics engine would address the current mechanical simplifications. These directions are feasible within the existing architecture; the knowledge base modularity that enables cross-cancer generalization also provides the entry point for richer biological content.

### STAR Methods Resource Availability Lead Contact

Further information and requests for resources should be directed to and will be fulfilled by the lead contact, Lianghua Wang.

### Materials Availability

This study did not generate new reagents or biological materials.

### Data and Code Availability

- All simulation source code, knowledge base YAML files, and analysis scripts are available at https://github.com/dawnmengsjtu/cellswarm.
- Simulation output data and intermediate results have been deposited at Zenodo (DOI will be provided upon publication).
- The TNBC single-cell reference dataset is publicly available at GEO: GSE176078 (Wu et al. 2021).
- No original code beyond that described above was generated for this study.

## Method Details

### Cell Agent Architecture

Each cell in CELLSWARM is modeled as an autonomous agent with four architectural layers.

#### Persistent state

Every agent maintains a mutable state vector comprising: cell_type (one of CD8^+^ T, Tumor, Treg, Macrophage, NK, B cell), two-dimensional grid position, energy (consumed by actions and replenished by glucose uptake), activation level, exhaustion level, proliferation rate, immune evasion score, suppressive activity, and polarization state. These scalar attributes are updated deterministically each simulation step before the decision phase.

#### Pathway state

Each agent carries a 14-dimensional signaling vector where every pathway takes a continuous value in [0,1]. The pathways are organized into five functional modules: immune activation (TCR, CD28), immune checkpoints (PD-1, CTLA-4), metabolism (mTOR, AMPK, HIF-1α), cytokine signaling (IFN-γ/JAK-STAT1, IL-2/JAK-STAT5, TGF-β/SMAD, NF-κB), proliferation and cell death (PI3K/AKT, MAPK/ERK, caspase). Pathway values are computed each step via a sigmoid activation function σ(x)=1/(1+e^-x^) applied to a weighted sum of environmental signals and internal state variables. For example, in a CD8^+^ T cell: TCR=σ(0.5·tumor_nearby+0.3·activation) and $\text{PD-1} = \sigma(1.2 \cdot \text{PD\text{-}L1\_local} + 0.5 \cdot \text{exhaustion})$. Weights are cell-type-specific and defined in the Pathway Knowledge Base.

#### Cell memory

Each agent maintains a two-tier memory stream: 5 short-term slots storing the most recent simulation steps (action taken, outcome, local environment snapshot) and 20 long-term slots reserved for salient events (e.g., first antigen encounter, onset of exhaustion, successful kill). Long-term memories are retained across the full simulation and provide episodic context to the LLM decision core in agent mode.

#### Cell cycle

Proliferation-competent cells follow a five-phase cycle: G0 (quiescent) ➔ G1 (8 h) ➔ S (6 h) ➔ G2 (4 h) ➔ M (1 h) ➔ G0, with each simulation step corresponding to approximately 1 h. Cells may also transition to apoptosis (triggered when caspase >0.6) or senescence (after exceeding a cell-type-specific division count). Division at M phase produces a daughter cell placed in an adjacent empty grid position.

### Knowledge Bases

CELLSWARM draws on five structured knowledge bases, all stored as human-readable YAML files.

*(1) Cancer Atlas* (6 files). Each file specifies a cancer type—TNBC, CRC-MSI-H, CRC-MSS, Melanoma, NSCLC, and Ovarian—and contains cell-type proportions, tumor growth rates, and immune infiltration parameters. TNBC parameters were derived from the GSE176078 single-cell RNA-seq dataset (Wu et al. 2021); parameters for the remaining five cancer types were curated from the tumor microenvironment classification of Bagaev et al. (Bagaev et al. 2021).
*(2) Drug Library* (22 monotherapies + 5 combinations). Each drug entry specifies: mechanism of action, target pathway(s), environment-level effects (e.g., signal field clearance), cell-level effects (e.g., pathway value modifiers), and cancer-specific dose modifiers. The library covers approved and investigational agents including pembrolizumab, nivolumab, atezolizumab, ipilimumab, and galunisertib.
*(3) Pathway KB* (15 files). Organized into five categories: immune activation (TCR signaling, antigen presentation), immune checkpoints (PD-1/PD-L1, CTLA-4/CD28, TIGIT/CD226), cytokine signaling (IFN-γ/JAK-STAT, IL-2/STAT5, TGF-β/SMAD, TNF/NF-κB), proliferation (PI3K/AKT/mTOR, RAS/MAPK, Wnt/β-catenin), and cell death (extrinsic apoptosis, intrinsic apoptosis, ferroptosis). Multiple KB files may map to the same signaling dimension (e.g., extrinsic and intrinsic apoptosis both feed into the caspase pathway value), yielding 15 knowledge base entries for 14 signaling dimensions. Each file defines upstream activators, downstream targets, and cross-talk edges used to construct the per-cell-type sigmoid weight matrices.
*(4) Perturbation Atlas* (6 files, one per cell type). Each file maps gene names to affected pathways and effect sizes: PDCD1 ➔ PD-1 (set to 0), CTLA4 ➔ CTLA-4 (set to 0), TGFB1 ➔ TGF-β/SMAD (set to 0), TP53 ➔ caspase (X0.5), IFNG ➔ IFN-γ/JAK-STAT1 (set to 0), BRCA1 ➔ caspase (X0.7), and IL2 ➔ IL-2/JAK-STAT5 (set to 0).
*(5) TME Parameters* (6 cancer-specific + 1 shared default). These files specify grid dimensions, diffusion coefficients for each signal field, signal decay rates (λ), and blood vessel positions for nutrient supply.

### Simulation Environment

The tumor microenvironment is modeled on a two-dimensional square grid whose size is dynamically adjusted to maintain a target cell density of 0.2 cells per grid point. Six diffusible signal fields are maintained: O_2_, glucose, IFN-γ, IL-2, TGF-β, and PD-L1. Signal dynamics follow a reaction-diffusion equation discretized via a 5-point Laplacian finite-difference stencil:

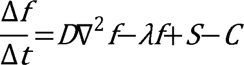

where D is the diffusion coefficient, λ the decay rate, S the local secretion, and C the local consumption. Tumor cells consume O_2_ at 0.002 per step; immune cells consume at 0.001 per step. Secretion rates are action-dependent: a CD8^+^ T cell attack deposits IFN-γ at +0.05; tumor cells secrete PD-L1 at +0.02Ximmune_evasion; Tregs secrete TGF-β at +0.04Xsuppressive_activity. Blood vessels at predefined grid positions supply O_2_=0.08 and glucose =5.0 each step. Immune cells exhibit chemotaxis, biasing migration toward the nearest tumor cell.

Simulations were initialized with 500 cells (200 tumor, 120 CD8^+^ T, 80 macrophage, 40 NK, 30 Treg, 30 B cell) and run for 30 time steps unless otherwise noted.

### Decision Modes

Three decision-making modes were compared, all operating on identical environment dynamics and pathway computations.

#### Agent mode

An LLM is queried to select each cell’s action. To optimize computational cost, LLM calls are gated by a complexity score defined as the fraction of pathways with activation >0.4 plus the variance across all 14 pathway values. Cells with complexity :::0.3 use a simplified rule-based decision pathway, as their low pathway variance indicates a microenvironmental context that does not require full LLM reasoning. This efficiency optimization reduces API calls by approximately 40% without affecting simulation fidelity (see ablation in Result 5). For qualifying cells, the LLM prompt comprises the cell’s persistent state, all 14 pathway values, memory context (short- and long-term), relevant KB entries, and a snapshot of the local 5X5 environment. The model returns a structured JSON object specifying action (one of: proliferate, migrate, attack, secrete, apoptose, quiesce), params (action-specific parameters), and secretion (cytokine outputs). Calls are batched and dispatched via a thread pool for concurrent execution.

#### Rules mode

Decisions are made by deterministic if-else rules applied to the same pathway values. For CD8^+^ T cells: net activation = TCR + CD28 - PD-1 - CTLA-4; if >0.5, attack; otherwise quiesce or migrate. For tumor cells: MAPK/ERK >0.5➔ proliferate; caspase >0.6➔ apoptosis; HIF-1α >0.5➔ migrate toward higher O_2_.

#### Random mode

Actions are selected uniformly at random from the available action set, serving as a null baseline.

### Combat Resolution

When an immune cell selects the attack action, it searches for tumor targets within Manhattan distance :::5. The kill probability is computed as:

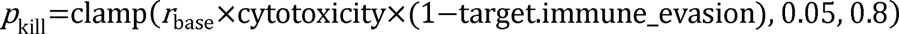

where r_base_=0.3. A Bernoulli draw determines whether the target is eliminated. Successful kills increment the attacker’s activation and deposit IFN-γ locally; failed attacks increment exhaustion.

### Treatment Simulation

Drug treatments are applied at either step 5 (early) or step 15 (late). Drug effects operate at two levels. First, *environment effects* modify signal fields directly (e.g., anti-PD-1 clears the PD-L1 field by a fraction proportional to drug strength). Second, *pathway effects* are applied after compute_pathways(): for anti-PD-1, the PD-1 pathway value is scaled as PD-1 +- PD-1 X(1-strength)^2^. Drug parameters are read from the Drug Library KB when available, with hard-coded fallback values otherwise. Three monotherapies were systematically evaluated: anti-PD-1, anti-CTLA-4, and anti-TGF-β.

### Perturbation Simulation

Seven single-gene knockouts were simulated: *PDCD1*, *CTLA4*, *IFNG*, *TGFB1*, *TP53*, *BRCA1*, and *IL2*. Perturbations are configured at initialization and enforced every step by calling_apply_perturbations() after pathway computation, which clamps or scales the mapped pathway value according to the Perturbation Atlas. Knockouts fall into two categories: *direct* KOs (*PDCD1*, *CTLA4*) alter pathways that appear explicitly in rule-based decision thresholds (e.g., PD-1 and CTLA-4 in the CD8^+^ T cell net-activation formula), and *indirect* KOs (*IFNG*, *TGFB1*, *TP53*, *BRCA1*, *IL2*) affect pathways outside the rule decision boundary, making their downstream effects accessible only to LLM-driven agents that can reason over the full pathway context.

### Phenocopy Analysis

To assess whether genetic perturbations recapitulate pharmacological interventions, we compared the final cell-type proportion vectors from knockout simulations with those from the corresponding drug treatments. Similarity was quantified by Pearson correlation across the six cell-type fractions. Observed correlations were: *PDCD1* KO vs. anti-PD-1 (r=0.86), *CTLA4* KO vs. anti-CTLA-4 (r=0.56), and *TGFB1* KO vs. anti-TGF-β (r=0.31), consistent with the expectation that direct-pathway knockouts produce stronger phenocopies than indirect ones.

### Ablation Experiments

Each ablation removed one knowledge base component (Cancer Atlas, Drug Library, Pathway KB, Perturbation Atlas, or TME Parameters) while keeping the remaining four intact.

Simulations were run with the DeepSeek backbone and 3 independent seeds per condition. Performance was assessed by JS divergence to the reference cell-type distribution.

### Computational Resources

Eight LLM backends were benchmarked: DeepSeek, GLM-4-Flash, Qwen-Turbo, Qwen-Plus, Qwen-Max, and Kimi-K2.5, plus the non-LLM Rules and Random baselines. A single 30-step simulation with 500 initial cells consumed approximately 2.8 M tokens in agent mode. Wall-clock runtime ranged from 8 to 50 minutes depending on the model backend. All simulations were executed on a single Apple MacBook Air (M1, 16 GB RAM) without GPU acceleration.

### Quantification and Statistical Analysis

Four metrics were used to evaluate simulation outcomes:

- **JS divergence**: Jensen–Shannon divergence between the simulated and reference (GSE176078; Wu et al. (2021)) cell-type proportion vectors, measuring compositional fidelity.
- **Tumor ratio**: Final tumor cell count divided by initial tumor cell count, quantifying net tumor growth or regression.
- **Shannon diversity**: H^1^=- ∑_i_ p_i_ lnp_i_ computed over the six cell types, capturing ecosystem heterogeneity.
- **CV**: Coefficient of variation (standard deviation / mean) of tumor ratio across seeds, measuring reproducibility.

Statistical comparisons used two-sided Welch’s t-tests. No correction for multiple comparisons was applied, as comparisons were pre-planned and limited in number. Throughout, n denotes the number of independent simulations initialized with distinct random seeds.

## Acknowledgments

This work was supported by the National Natural Science Foundation of China (NSFC, No. 82173732), the AI for Science Project (Grant No. RGZD002) of the Shanghai Municipal Education Commission, the Yizhang Talent Plan (JCYZRC-D-037), and the Basic Medical Research Program of Naval Medical University (2025QN004).

